# Identifying cellular-to-phenotype associations by elucidating hierarchical relationships in high-dimensional cytometry data

**DOI:** 10.1101/2021.07.08.451609

**Authors:** Adam Chan, Wei Jiang, Emily Blyth, Jean Yang, Ellis Patrick

## Abstract

High-throughput single cell technologies hold the promise of discovering novel cellular relationships with disease. However, analytical workflows constructed for these technologies to associate cell proportions with disease often employ unsupervised clustering techniques that overlook the valuable hierarchical structures that have been used to define cell types. We present treekoR, a framework that empirically recapitulates these structures, facilitating multiple quantifications and comparisons of cell type proportions. Our results from twelve case studies reinforce the importance of quantifying proportions relative to parent populations in the analyses of cytometry data — as failing to do so can lead to missing important biological insights.

## Introduction

High-parameter cytometry assays have provided biomedical scientists with an unprecedented detail of the cellular heterogeneity of patient samples. Flow and mass cytometers are able to characterise cells by measuring up to fifty extracellular antigens^1^, with single-cell sequencing platforms able to measure thousands of intracellular RNA molecules^2^. Unfortunately, this ground-breaking capacity to characterise cells to this depth has provided a computational challenge for bioinformaticians to efficiently glean meaningful information from the deluge of single-cell data. Given that most novel analytical methods neglect the hierarchical relationships in single-cell data, there exists an opportunity to use these relationships to identify robust and interpretable associations between cell subsets and patient clinical end points or *ex vivo* interventions.

To compare the abundance of cell subsets between samples, there has been a decades-long legacy of either quantifying a cell type as the proportion relative to all cells in a sample (*%total*), or, as the proportion relative to a parent population (*%parent*)^3–5^. The latter of these quantifications is derived naturally from the way that cell subsets have traditionally been annotated via a process called sequential manual gating^6^ - where 2D scatter plots are drawn using certain markers and gated to identify cell populations in a sequential manner. For example, regulatory T cells (Tregs) could be identified by first gating out CD3+ and CD4+ cells to identify CD4+ T cells and then further gating on CD25lo and CD127+ to isolate the CD4+ Tregs^7^. This gating strategy naturally lends itself to quantification of cell types relative to their parent lymphocyte populations. These quantifications are robust to changes in unrelated subsets. The main drawbacks of this method however are its reliance on time-consuming manual gating, which has become impractical for high-parameter assays^8^, and the substantial reliance on expert knowledge which may bias analysis towards known and expected relationships.

As an alternative cell type identification strategy to manual gating, unsupervised clustering of cells has been used to circumvent the challenges of sequentially gating high-dimensional cytometry data. These automated methods are able to stratify cell subsets without necessarily having a predetermined hypothesis or sequential gating strategy. Many methods, including SPADE^9^, Citrus^10^, FlowSOM^11^, Phenograph^12^, SC3^13^ and scClus^t14^ have been utilised frequently in the analysis of high-dimensional cytometry data to identify cell populations. Whilst they have significantly improved the efficiency in which scientists can analyse these datasets, typical analyses employing these methods only explore the changes in cell types as a %total, neglecting the complex hierarchical proportions inherent in single cell data. In other words, these methods fail to measure cell types as a %parent, which cytometry analysts have traditionally used in manual gating workflows.

A number of unsupervised clustering methods and data-driven workflows have been developed to explore the hierarchical nature of cytometry data. SPADE and FlowSOM utilise minimal spanning trees over clustering as a visualisation tool. Citrus employs hierarchical clustering and regularised supervised learning algorithms to identify stratifying populations of cells on each level of aggregation. The method treeclimbR^15^, aims to pinpoint an ideal resolution of cell populations via a hierarchical tree. Although these methods acknowledge the importance of visualizing the hierarchical aspect of single cell cytometry data, they do not typically incorporate such information in their association analysis. That is, they do not by default quantify the abundance of cell types as a %parent and test if these compositions are associated with a treatment or phenotype of interest.

To this end we have developed treekoR, a novel framework that makes use of cell type identification from unsupervised clustering techniques whilst acknowledging the hierarchical nature of single cell cytometry data to discover robust and interpretable associations between cell subsets and patient outcomes. TreekoR achieves this by (1) algorithmically deriving the hierarchy of cell type clusters, followed by (2) incorporating this hierarchical information via measuring the %parent for each cell type. These derived proportions can then be used in significance testing and classification models to determine associations with clinical outcomes. Further to this, treekoR provides a general framework that is flexible to the clustering approach, hierarchical aggregation method, and type of significance testing used. This framework allows analysts to generate insight from the complex hierarchical relationships present in single cell cytometry data, which are often overlooked with existing automated clustering methods.

## Results

### treekoR algorithmically derives cell type hierarchies to quantify %parent

We present treekoR, an analytical framework that recognizes and incorporates the hierarchical relationships inherent in cytometry data. The treekoR package is implemented in R and uses an automated workflow to identify cellular associations with a patient outcome through five steps (**Figure 1**): (1) cluster the data using an automated method, (2) aggregate clusters into a tree using a hierarchical clustering algorithm, (3) calculate the %total (the proportion of a cell type relative to all cells in a sample), and %parent (the proportion of a cell type relative to a parent population of cells, in this case the cells in the parent node) of cells in each node in the tree, (4) perform significance tests using both of these proportions against a clinical end point, and (5) visualise the significance results on the tree. The %parent calculated by treekoR aims to emulate the proportions naturally derived when using sequential manual gating, which are not typically calculated in workflows exclusively using unsupervised clustering methods. Our comparative procedure uncovers important associations with a clinical end point of interest by visualising both quantifications of cell type proportions derived from the data. Further details are provided under Methods.

**Fig 1:**
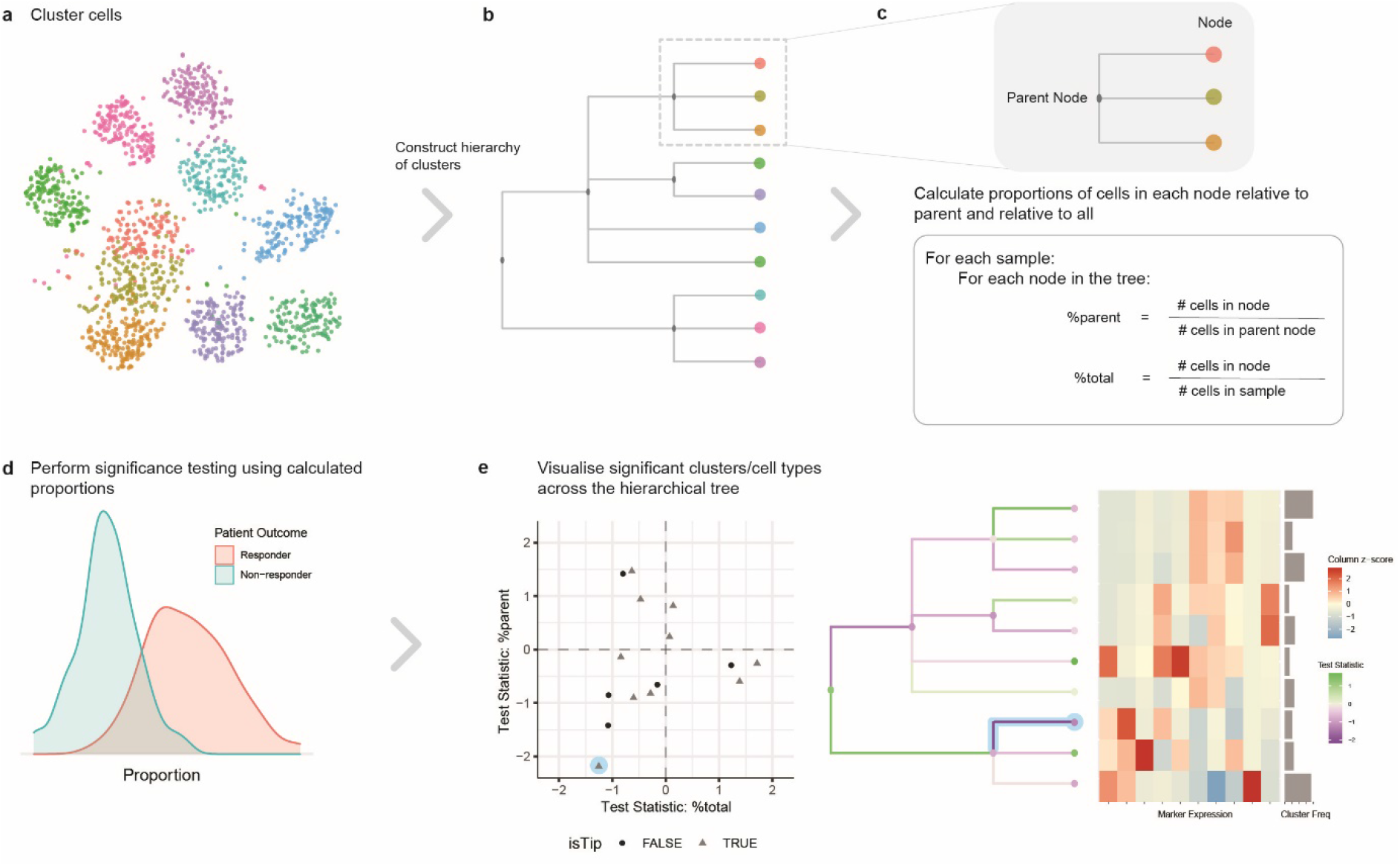
treekoR helps to extract insight from cytometry data through deriving a hierarchy of cell clusters and measuring proportions to parent. **a.** An example t-SNE plot showing clustering of single cell data. **b.** Hierarchical tree constructed using HOPACH algorithm on the cluster median marker expressions. **c.** Definition of proportions to parent and proportions to all defined according to the organisation of the hierarchical tree. **d.** Significance testing is performed using both types of proportions calculated, testing for difference between the patient clinical endpoint of interest. **e.** Visualisation of the significance testing results. On the left, a scatterplot of each node in the hierarchical tree with the test statistic calculated using the %total (x-axis) vs. the test statistic calculated using the %parent (y-axis). On the right of the scatterplot, the hierarchical tree is coloured with the test statistics: the nodes coloured by the test statistic using %total and the branches of the nodes coloured by the test statistic using %parent. An example of a corresponding node between the two graphs is highlighted in blue. The heatmap plots the median marker expression of the leaf nodes to assist in identification of the corresponding cell clusters.

### treekoR generates biological insight exclusive to %parent in example cytometry datasets

We illustrate the ability of treekoR to generate additional biological insight by applying the framework to a CyTOF study of latent Cytomegalovirus (CMV)^16^. After clustering cells into one hundred cell subsets, quantifying the %total and %parent for each, and testing for associations between CMV positivity and %total or %parent (**Figure 2a**); we observed a reduction in CD4+ Tem cells in CMV positive patients using %parent (p=6.1× *10*^−*5*^, FDR=3.33× *10*^−*3*^), yet no association was observed using %total (p=0.9, FDR=0.99). The higher proportion of CD4+Tem relative to its parent cluster (CD4+Tem and CD4+Tcm) in CMV negative patients as compared to CMV positive patients is in keeping with known effector memory cell function in cytokine secretion and viral clearance. Similarly, observed a nominally significant negative association between CMV positivity and CD8+ CD127-Tem cells using %parent (p=1.5× *10*^−*3*^, FDR=3.5× *10*^−*2*^), but not with %total (p=0.26, FDR=0.69) (**Figure 2b**). This lower proportion of CD8+ CD127-Tem cells relative to its parent (CD8+ CD127- and CD8+ CD127+ Tem) in CMV positive patients as compared with CMV negative patients suggests a role for differential CD127 expression in chronic/persistent infection. Together, this suggests that if the %parent of these cell types had not been measured, we would have been unable to discover the cellular relationships between CD4+ Tem and CD8+ CD127-Tem with CMV infection.

**Fig 2:**
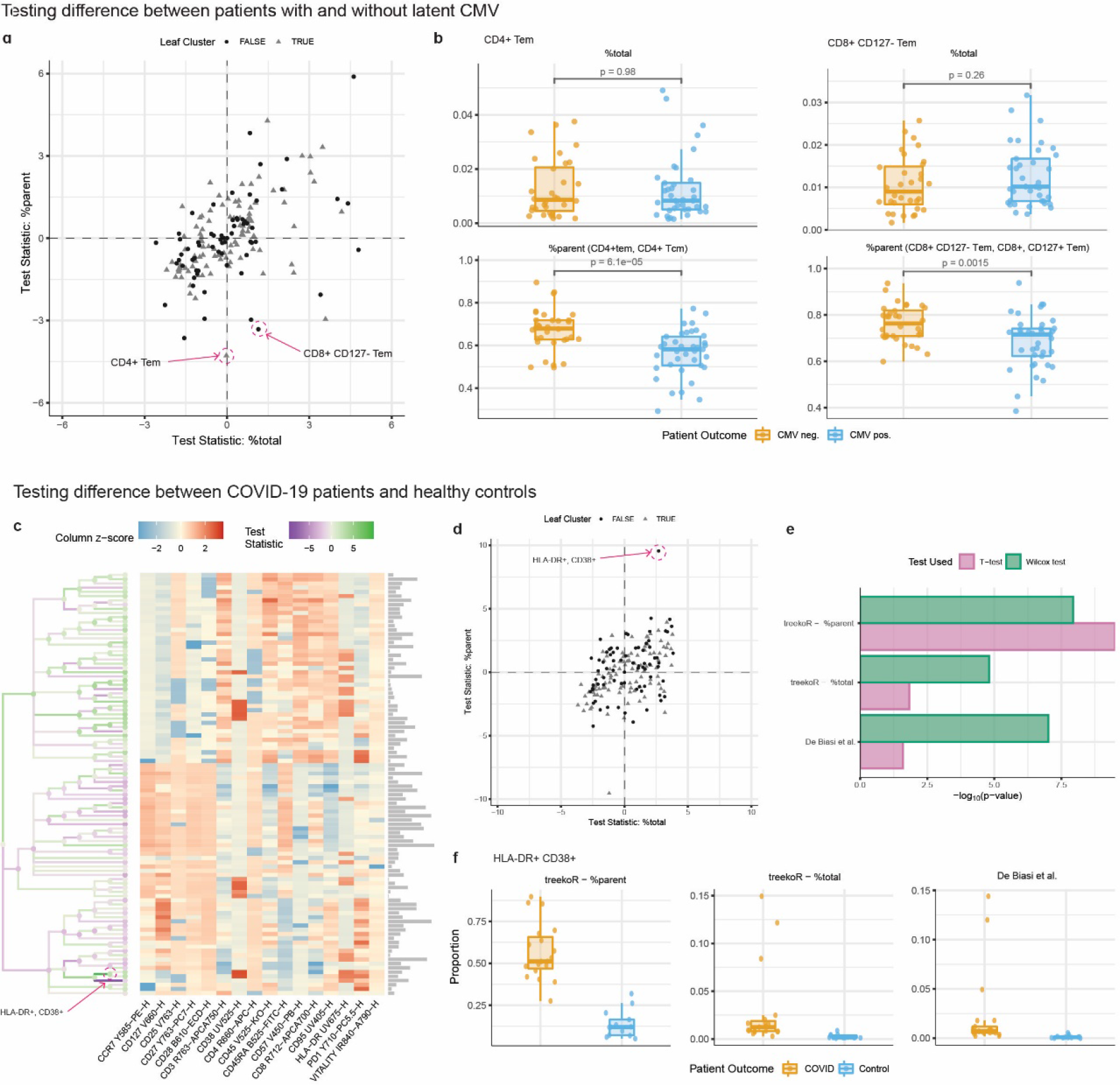
Measuring %parent can provide additional insight over %total. **a.** Scatterplot of test statistics with the cell clusters in differentiating between latent CMV infection patients. Highlighted clusters are significant using %parent, whilst not significant using %total. **b.** Comparative boxplot of the proportions of highlighted cell clusters, between patients with CMV and without CMV, with the %total (upper panel) and %parent (lower panel). **c**. A heatmap generated using treekoR on a CD8+ T cell compartment to predict healthy vs COVID-19, containing a hierarchical tree of cell clusters coloured by the test statistic using the corresponding %total (nodes) and %parent (branches). The heatmap is coloured by the scaled cluster median expression values characterise leaf nodes in the tree. **d.** Scatterplot of test statistics of each cell cluster with test statistic from using %total (x-axis) vs. test statistic from %parent (y-axis). The HLA-DR+ CD38+ cluster highlighted has a larger test statistic when differentiating between COVID-19 patients and healthy control using %parent than %total. **e.** Comparison of −log10 of p-values of a HLA-DR+ CD38+ subset for %total, %parent and manually gated proportions from De Biasi et al. from a t-test (pink) and Wilcoxon test (green). **f.** Comparative boxplot of a HLA-DR+ CD38+ subset, with the %total (upper left panel), %parent (lower panel), and manually gated proportions (upper right panel) between COVID-19 and healthy patients

When applied to a flow cytometry dataset profiling CD8+ T Cells in COVID-19 patients and healthy controls^4^, treekoR highlighted a highly activated HLA-DR+ CD38+ CD8+ T cell subset whose %parent provided a more robust association with COVID-19 response than its %total. After applying FlowSOM to cluster cell types (**Figure 2c**), we discovered a HLA-DR+ CD38+ CD8+ T cell whose %parent is greater in COVID-19 patients than healthy controls (p=3.19× *10*^−*10*^, FDR=2.76× *10*^−*8*^) (**Figure 2d**). However, this population only appeared marginally associated with COVID-19 response using %total (p=1.49× *10*^−*2*^, FDR=*7*.*6* × *10*^−*2*^). In contrast, De Biasi et al., had reported a manually gated HLA-DR+ CD38+ CD8+ T cell population changing when using %total (p=9.70× *10*^−*8*^). The difference in conclusion between using %total from FlowSOM and the manually gated population from De Biasi et al. is solely attributed to our use of a t-test and De Biasi et al.’s use of the Wilcoxon rank sum test (**Figure 2e**), which is robust to the outliers observed in the %total quantification (**Figure 2f**). When a Wilcoxon rank sum test is used on our %total (p=1.55× *10*^−*5*^, FDR=1.12× *10*^−*3*^) and %parent (p=1.15× *10*^−*8*^, FDR=9.96× *10*^−*7*^) the association is also observed, but not observed when a t-test is used on De Biasi et al.’s manually gated population (p=2.57× *10*^−*2*^). The presence of this association in treekoR’s %parent regardless of the significance test used illustrates that quantifying the proportion of HLA-DR+ CD38+ to a parent population (HLA-DR+ CD38+ and HLA-DR+ CD38-) can adjust for large fluctuations in cell type compositions and allow subtle changes in proportion to be robustly quantified. Across both the COVID-19 and CMV case studies we highlight two perspectives of cell type proportions, %total and %parent, which offer biological information that may be potentially missed if only one was measured.

### The %parent of cell types yields strong associations with clinical outcomes across several datasets in our benchmark

We observed a greater discrimination between binary outcomes through quantifying proportions as %parent than %total in several datasets. We compared twelve case studies consisting of seven CyTOF datasets, four flow cytometry datasets and a single-cell RNA sequencing (scRNA-seq) dataset (**Table 1**). Further, we also used two hierarchical clustering algorithms, HOPACH^17^ and average-linkage hierarchical clustering, with both generating different estimates of %parent (**Supplementary Figure 1**). After testing for differences in cell type proportions between the patient conditions, we compared the ordered negative log p-values of each cell population from using %total against the ordered negative log p-values from using %parent (**Figure 3**). Across all twelve case studies, we were able to determine whether performing significance testing using %parent provided comparatively stronger associations with the patient outcome than %total - evident in instances where points conspicuously lay above the dashed identity line. Across half of the investigated datasets, in particular CMV^16^ and Age Chronic^18^, the cell type proportion with highest significance was obtained from measuring its %parent. Further to this, the choice of hierarchical aggregation techniques produced variations in clinical association, suggesting that using different cell type trees can help analysts uncover a wider scope of associations. The benchmark exemplifies the importance of measuring both %parent and %total so as not to miss pertinent clinical associations.

**Table 1:**
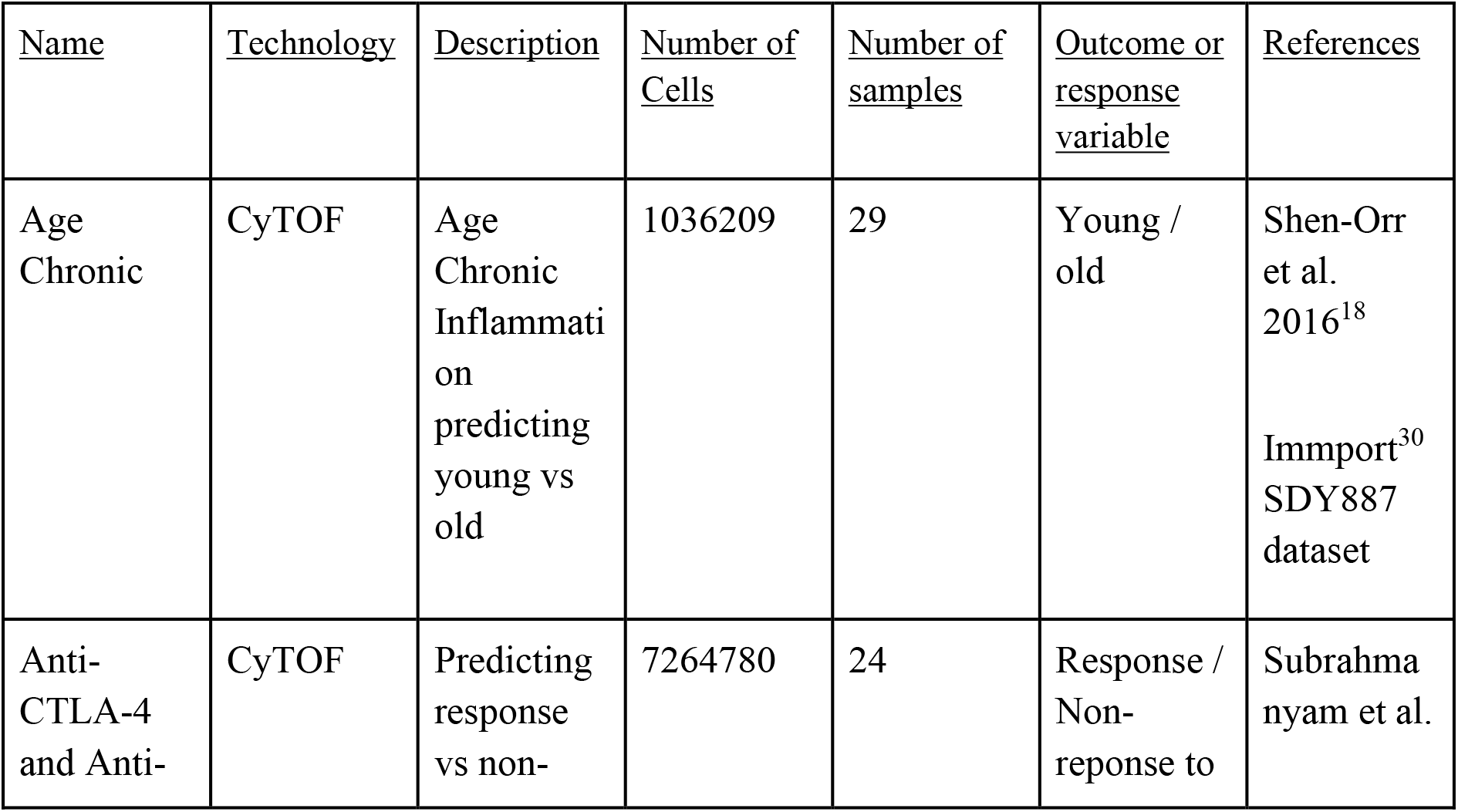
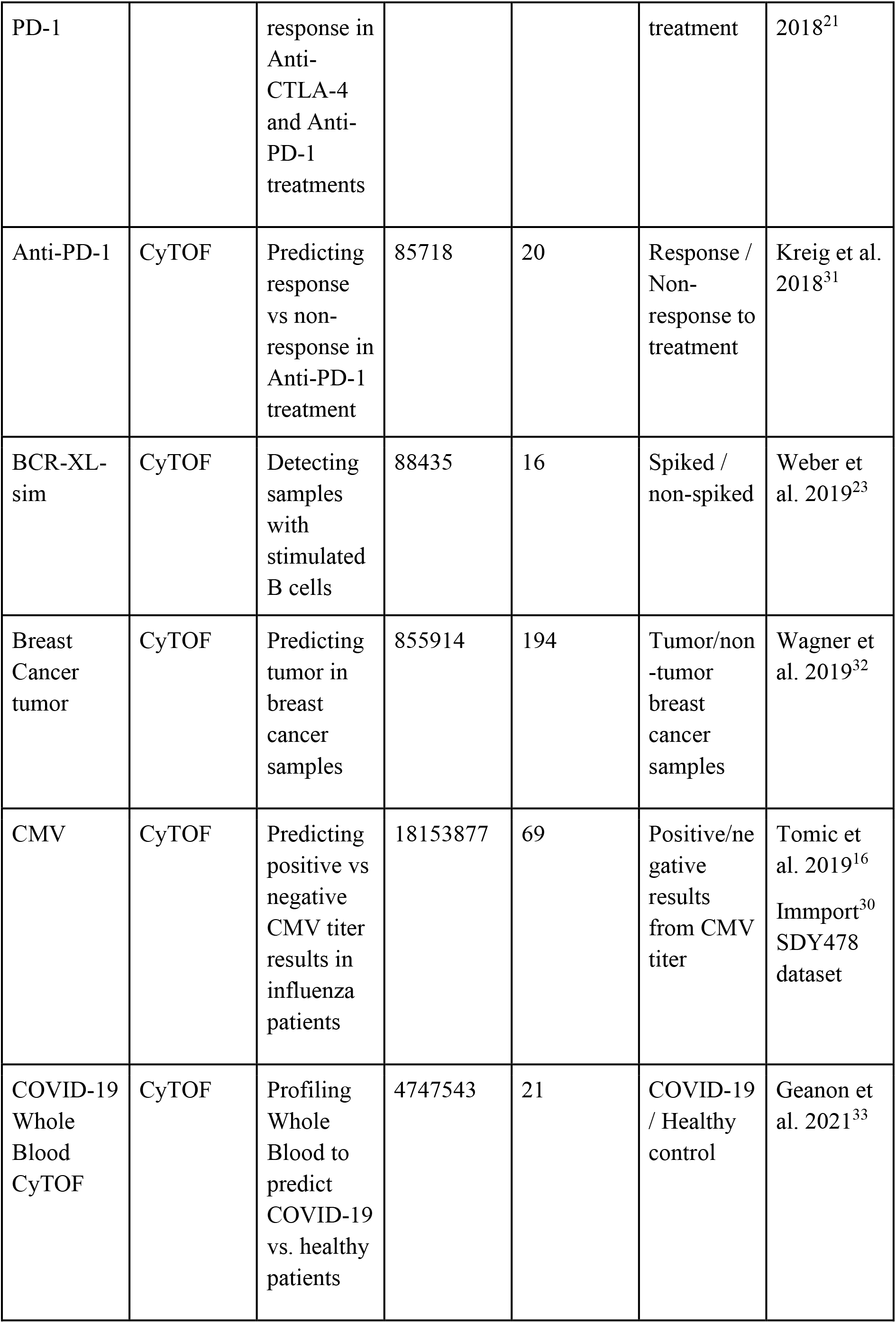
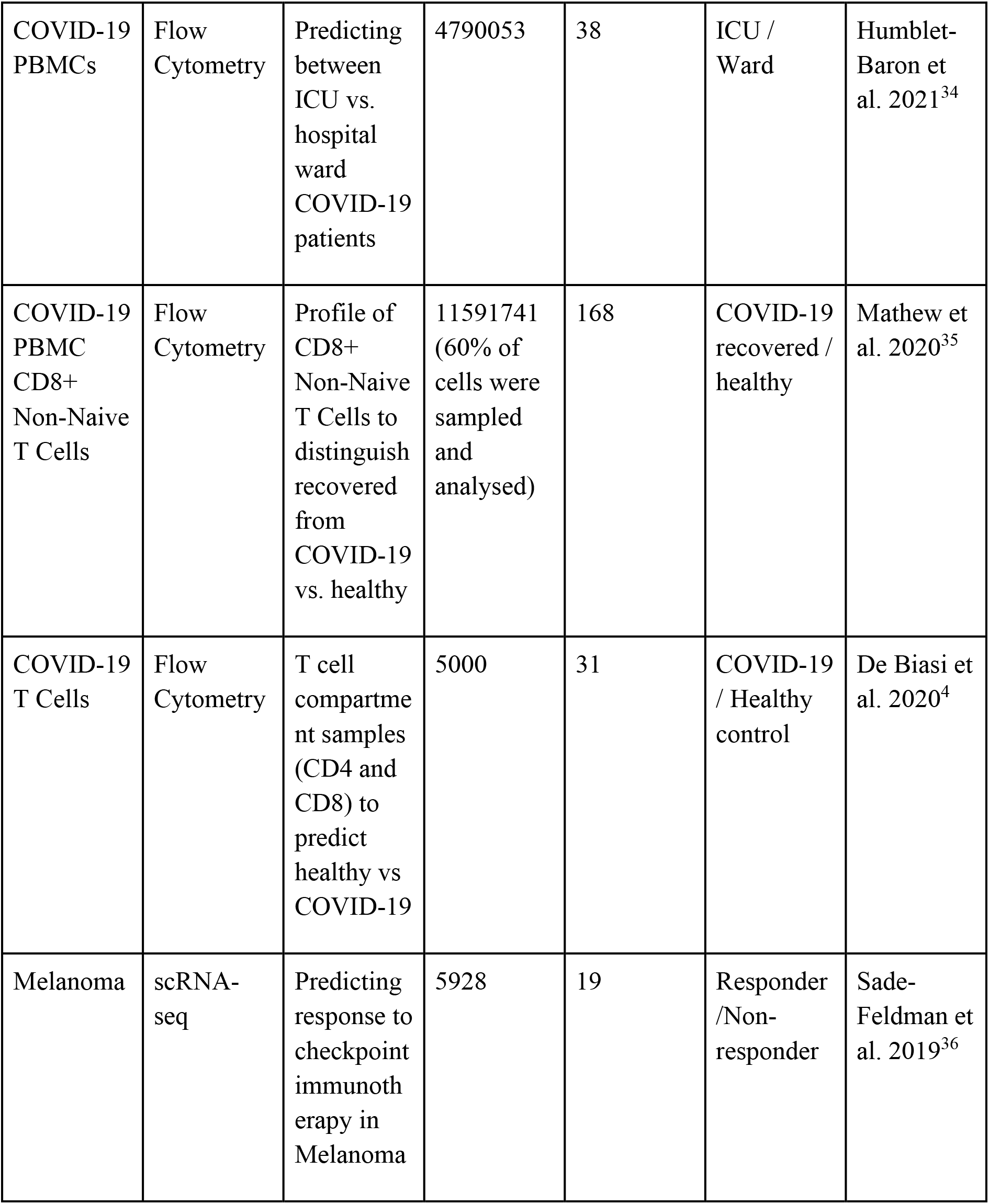
Benchmark datasets. Eleven published datasets were used to compare %total and %parent in significance testing and classification using the treekoR workflow. “Name” is used to refer to each dataset throughout the manuscript.

**Fig 3:**
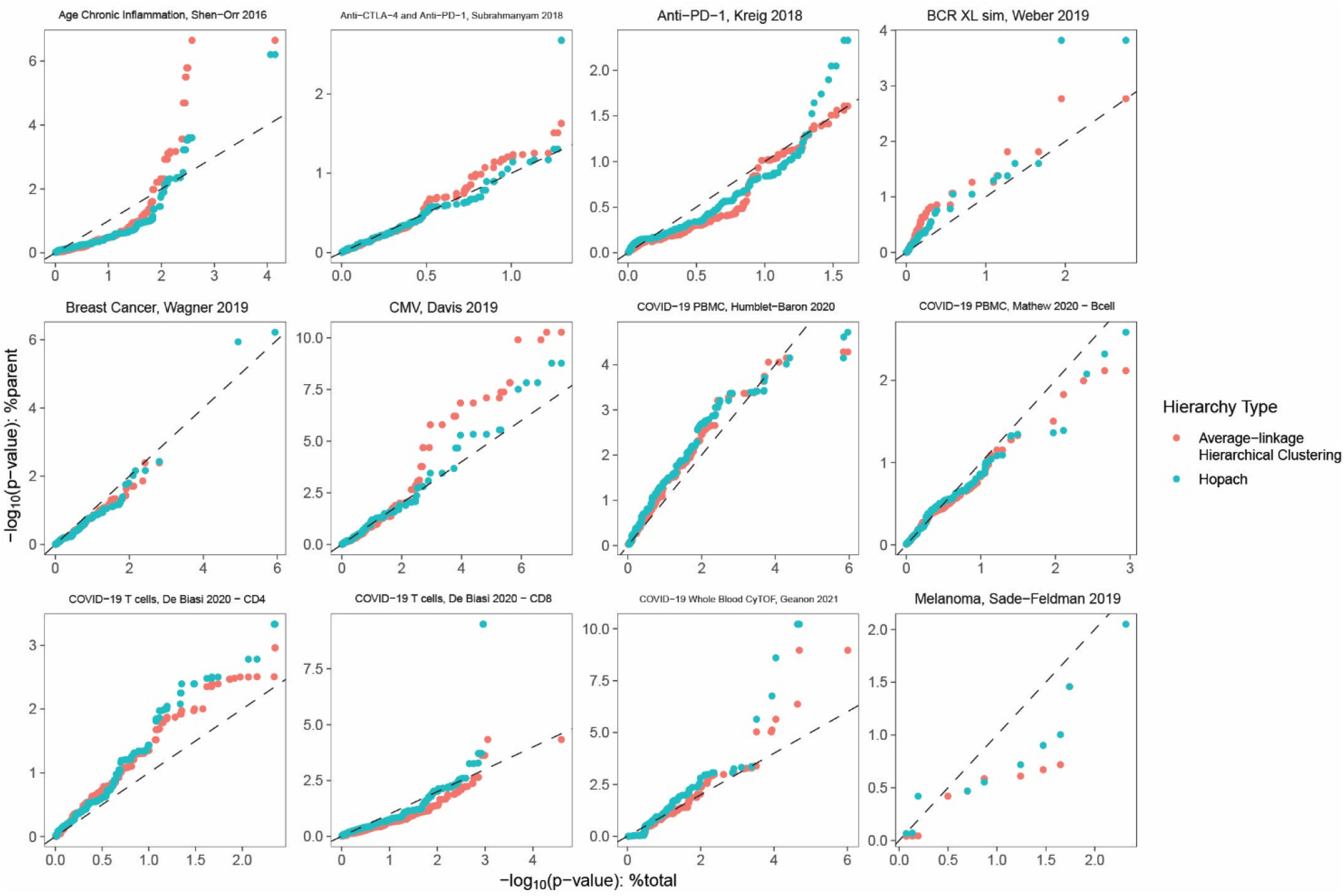
treekoR provides stronger associations with patient clinical outcomes. Cell clusters, constructed using both average-linkage hierarchical clustering and HOPACH, were tested between patient conditions using %total and %parent. Q-Q plots were plotted for each dataset by plotting the ordered negative log p-values using %total (x-axis) vs. using %parent (y-axis).

### Multivariate classification of clinical outcomes in cytometry data can be improved by measuring %parent

High-dimensional single cell data have been used to construct models to classify patients to help scientists discover and understand associations with a clinical outcome^19–22^. We evaluated classification performance using either %total or %parent as feature sets in several datasets with binary outcomes (e.g. responder vs. non-responder, COVID-19 vs. healthy control), to determine that the incorporation of %parent features in multivariate classifications models can help improve patient classification. There were various differences in balanced accuracy between using %total and %parent (using either HOPACH or hierarchical clustering with average linkage) in each dataset (**Figure 4**). The datasets with the biggest increase in balanced accuracy by using %parent were the BCR-XL-sim data^23^ and Age Chronic data^18^. In the BCR-XL-sim semi-simulated dataset, we predicted which samples contained stimulated B cells. Using only %total as features produced a mean balanced accuracy of 59%, compared to 73% using %parent derived from HOPACH. In the Age Chronic CyTOF dataset, classifiers were constructed to discriminate between older and younger adults using their immune response signatures to influenza vaccination. Here, we show that using %parent (99%) also gives a higher mean balanced accuracy than %total (88%). These results support the notion that failing to measure %parent can sometimes mean neglecting important signals when trying to predict a patient’s clinical outcome in high-dimensional cytometry datasets.

**Fig 4:**
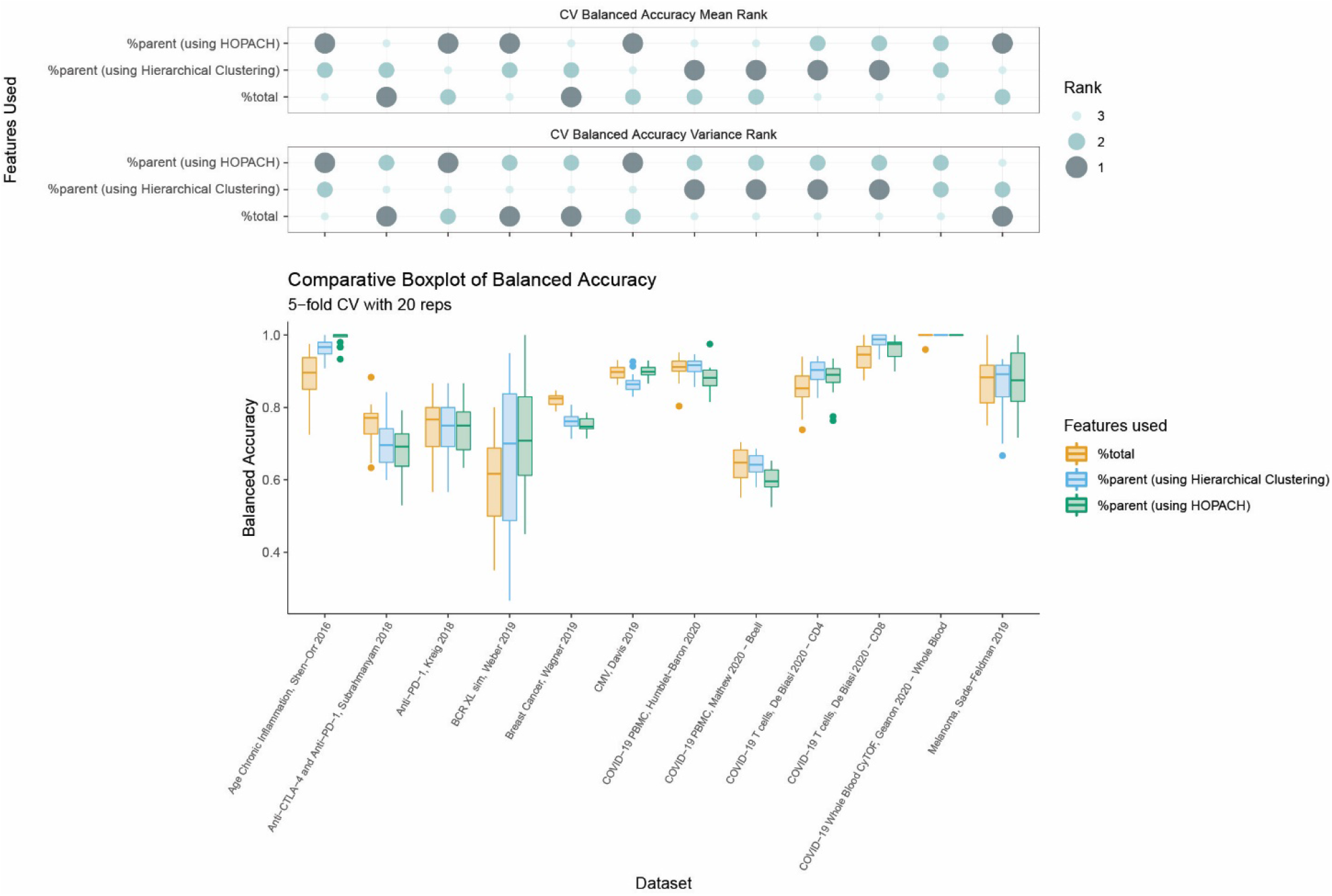
Measuring %parent offers improvements in patient classification performance. **a.** Comparative boxplots (lower panel) of balanced accuracy rates for each dataset and feature set: %total, %parent using average-linkage hierarchical clustering, and %parent using HOPACH. Values plotted are from a 5-fold CV with 20 repetitions, averaged across each repetition. The rank of each feature set within each dataset is shown in the bubble plot (upper panel), with rank 1 being the best (highest mean / lowest variance) and rank 3 being the worst (lowest mean / highest variance).

Quantifying multiple views of cell type proportions can provide greater insight into single cell cytometry data and patient clinical outcomes. In our classification benchmark, we compared the use of %total, %parent (using hierarchical clustering) and %parent (using HOPACH) cell type proportions. Exploring hierarchical representations via treekoR can help to elucidate a broader scope of %parent relationships that exist within cytometry data (**Supplementary Figure 1**). When each feature set was ranked using the mean and standard deviation of the balanced accuracy in each dataset (**Figure 4**), no single quantification of proportion performed the best for prediction of patient outcomes across all analysed cytometry datasets. The differences in rank however means that each type of proportion quantification provided a different perspective of the data. Depending on the dataset, one approach may provide a greater coverage of the signal present within the data through a higher balanced accuracy. This further supports the idea that proportions measured as %total should not be the only proportions measured in cytometry analysis workflows, particularly when searching for the most predictive features in distinguishing between patient clinical outcomes and understanding the complex relationships that exist. It is therefore imperative that proportions are quantified as both %parent and %total for the effective analysis of cytometry data, as it offers more thorough examination of this data.

## Discussion

In this paper, we examined several high-dimensional single cell datasets to demonstrate the importance of measuring both %parent and %total proportions, the use of %parent for classification and the consequences of using different hierarchical aggregation techniques to empirically derive cell type proportions. Overall we accentuated the importance of analysing high-dimensional cytometry data using ideas from both traditional manual gating and unsupervised clustering techniques, and provide a general framework, treekoR, which allows analysts to do so whilst overcoming key pitfalls of both approaches.

The treekoR framework allows scientists to select their own clustering algorithm for determination of cell types and hierarchical aggregation technique for the construction of cell type trees. Whilst there have been numerous comparisons of clustering methods of cytometry data^19,24–27^, there have not been as many comparisons of hierarchy construction techniques in the context of cell type hierarchies^9,11^. We show through the use of HOPACH and average-linkage hierarchical clustering that the choice of hierarchical aggregation technique can have noteworthy effects on downstream analysis, and suggest multiple other techniques that could also be used to produce distinct cell type trees. However no formal evaluation to determine the most ‘suitable’ technique was performed throughout our analyses. Since scientists have unique and personal workflows for hierarchically analysing cell types, there is significant room to explore what an appropriate cell type hierarchy might entail and determine a corresponding standard or measure which scientists can use to evaluate this. The definition for the most ‘suitable’ hierarchical aggregation technique, whether it is the technique which produces the most interpretable hierarchy or produces the %parent proportions most associated with a clinical outcome, has yet to be elaborated.

In treekoR we defined %parent as the proportion of a cell type relative to its direct parent in the cell type hierarchy. This proportion could be calculated using a broader parent (e.g. a higher ancestor) cell type in the hierarchy, which could lead to either a more interpretable and familiar cell type %parent or reduce the burden of multiple hypothesis testing. Since the scope of proportions to be calculated becomes much larger when numerous measurements of %parent for a single cell type are allowed, there exists a challenge in determining which %parent to calculate, particularly as the number of hypothesis tests increases. We do not currently address either of these points in our workflow. To overcome this challenge, a standard set of reference cell types can be determined to calculate %parent from. These reference cell types could be deduced in a semi-supervised fashion where analysts manually select them, or in a completely unsupervised manner by using a data-driven method (such as treeclimbR^15^). This would limit the number of proportions calculated and potentially provide more biologically relevant %parent.

Care is required in the comparison of statistical significance between the %total and %parent of a cell type. The derived p-values from significance testing inherently come from two distinct statistical hypotheses. Therefore the user should not conclude that one proportion is a better metric based solely on its p-value, or say that one proportion is more relevant than the other. Rather the %total and %parent provide two complementary views, both of which may be objective and biologically relevant. Depending on the datasets, one quantification of cell type proportions may provide a stronger association with a clinical outcome of interest, this nuance is important to note.

In summary, we present a framework that is general in nature, allowing scientists to choose algorithms appropriate to their dataset to glean more information than typical analyses. It is our broader intention to emphasise the importance of measuring %parent in the analysis of cytometry data - and that these hierarchical proportions should not be overlooked as researchers move towards more efficient and automated approaches of analysis. As high-dimensional cytometry data become more ubiquitous in helping scientists understand the underlying biological process behind patient diseases, such as influenza and COVID-19, we envision that the implementation of treekoR will assist in unravelling the cell type heterogeneity present in these complex patient diseases.

## Methods

### Overview of treekoR

treekoR is performed in five steps: (i) cluster the data using an automated method, (ii) aggregate clusters into a tree using a hierarchical clustering algorithm, (iii) calculate the %total and %parent of cells in each node of the tree, (iv) perform significance tests using both of these proportions against a clinical end point, and (v) visualise the significance results on the tree. The steps are described in detail below, along with the parameters used in the analyses throughout the manuscript.

#### (i) Clustering

Unsupervised clustering was performed using the FlowSOM^11^ algorithm as part of the CATALYST^28^ package in R^29^, using a 10×10 grid. Cells are over-clustered to try to account for all cell types present within the data and to avoid missing rare cell populations (any superfluous clusters are then naturally aggregated in the hierarchical clustering step). For the datasets that were provided with previously analysed or manually gated cell types, those cell types were used instead of the FlowSOM clustering.

#### (ii) Construction of hierarchy

Following clustering of the data, the scaled median marker expression for each cluster was calculated and used to construct a hierarchical tree. Several hierarchical clustering techniques can be used in treekoR and are included in the R stats hclust function^29^. These include HOPACH and agglomerative hierarchical clustering using: average-linkage, Ward-linkage, single-linkage, complete-linkage and McQuitty. HOPACH allows for multiple children per node whilst other included methods only cater for two children per node. Throughout the analysis we used two main methods for hierarchical aggregation: HOPACH (with *K* = 5 maximum children per parent node) and average-linkage hierarchical clustering.

#### (iii) Calculation of proportions

After clustering, when a hierarchical tree of cell types has been established in the data, the proportions of these clusters are quantified. For each patient, the proportions of cells belonging to the clusters in each node of the tree are measured relative to their total number of the cells, referred to as %total. In addition, the clusters in each node of the tree are measured as a proportion of the cluster in the direct parent node of the tree, referred to as %parent.

#### (iv) Significance testing

For each node in the hierarchical tree on the clusters, significance testing is then performed using a two sample t-test for equal means between the desired patient outcome using both the %total and %parent.

#### (v) Visualisation

The results of these proportions can be then visualised through a coloured tree plotted next to a corresponding heatmap. The heatmap displays the median scaled marker expressions of each cluster to help understand what cell type each cluster may represent, and the tree not only reveals how clusters have been hierarchically aggregated, but is coloured on each node by the test statistic obtained when testing using %total of that node, with the branch connecting the child to the parent coloured by the test statistic obtained when testing using the %parent of the child node.

### Benchmark data and data processing

The twelve benchmarking datasets consist of seven CyTOF, four flow cytometry and one single-cell RNA-seq dataset as shown in Table 1. In the flow cytometry datasets, COVID-19 T cells were counted as two datasets - CD4 and CD8 T cells.

### Data normalisation

For each of the cytometry datasets, we applied an arcsinh transformation with a co-factor of 5 on the expression values. The samples were then filtered to only include the patients with the clinical end points of interest. For analysis of the CMV dataset, 66.67% of cells were randomly subsampled and gated for live intact cells before transforming.

### Calculation of proportions

For each of the patients/samples, the proportions of each of the FlowSOM clusters or cell types were calculated as %total, as well as %parent from a HOPACH^17^ tree and an average-linkage hierarchical clustering tree. The %parent for each cluster in each sample is calculated as the (# cells in a cluster) / (# cells in a cluster + # cells in sibling clusters). The %total is calculate as (# cells in a cluster) / (# cells in sample).

### Hypothesis testing

For each of the cell types/clusters, a two-sample t-test was used to test if there was a significant difference in mean proportion between the binary clinical outcome of interest, using both %total and %parent. In our COVID-19 T cells and CMV case studies, p-value adjustment was performed using the FDR method, whilst p-value adjustment was not performed in the benchmark comparison.

### Classification

The %total and %parent proportions were then used as features separately, for sake of comparison, to predict the binary patient clinical end point. For each feature set and dataset combination, we trained a random forest (using mlr3^37^) with 500 trees in each iteration of a 5-fold cross validation with 20 repetitions. The balanced accuracy was measured in each iteration of the cross validation and used to compare predictive power between the feature sets.

All analysis was done in R^29^ version 4.0.3.

## Code availability

The code to run treekoR is available on Bioconductor (https://bioconductor.org/packages/release/bioc/html/treekoR.html) and code to reproduce the manuscript analysis from processed data has been shared on Github (https://github.com/adam2o1o/treekoR_analysis)

## Author Contributions

EP conceived and designed the study with input from JY. EP and AC led the treekoR method development and JY developed and guided the evaluation data analysis. AC curated the benchmarking data, implemented all data analytics and developed the R package with guidance from EP. WJ and EB contributed the biological interpretation of statistical findings. All authors wrote, read, reviewed the manuscript and approved the final version.

## Acknowledgements

The authors thank all their colleagues, particularly at The University of Sydney, School of Mathematics and Statistics, Charles Perkins Center, for their support and intellectual engagement. The following sources of funding for each author, and for the manuscript preparation, are gratefully acknowledged: Australian Research Council Discovery Project grant (DP170100654) to JYHY; Australian Research Council Discovery Early Career Researcher Award (DE200100944) funded by the Australian Government to EP; Australian Government Research Training Program (RTP) Scholarship to AC; PhD scholarship from the Haematology Society of Australia and New Zealand (HSANZ) and Leukaemia Foundation Australia to WJ. EB is a NSW Cancer Institute Post-Graduate Fellow and is supported by funding from NSW Ministry of Health, NSW Cancer Institute, Cancer Council of NSW, the Leukaemia Foundation of Australia and a research grant from MSD.

## Declarations

### Competing interests

The authors declare that they have no competing interests.

### Data Availability

All data analysed during this study are included in the published articles in **Table 1**. Benchmarking data generated in this study is available in the treekoR_analysis repository on GitHub, https://github.com/adam2o1o/treekoR_analysis.

## Supplementary

### Different hierarchical clustering methods uncover varying %parent relationships in cytometry data

In this section several tree structures derived from different hierarchical aggregation techniques are compared, in addition to a manual gating tree structure. The manner in which these ultimately result in capturing different signals from the data are demonstrated. Although no single representation may be the absolute correct one, exploring these different representations can begin to help analysts to uncover a broader scope of complex relationships that exist within cytometry data to discover the cellular heterogeneity between patient samples.

In our framework, the HOPACH algorithm was used to hierarchically aggregate the clusters into a tree. HOPACH is a clustering algorithm that was originally developed for gene expression data analysis, but has useful properties for the analysis of high-dimensional cytometry data. One advantage is the lack of restriction for splits to be binary, allowing up to 15 child nodes per node (**Supplementary Figure 1a**). In this example it can be observed that Pre-B cells, Mature B Cells and Immature B Cells fall under the same parent node, which, for example, would allow analysts to determine whether the compositional makeup of B cells plays any role in patient disease. Another advantage of HOPACH is the cluster collapsing step, which helps to alleviate any incorrect splitting of clusters. This helps prevent the tree from containing too many branches, which can reduce some correlations between proportions as our framework explores each of the parent-child relationships in the generated trees.

The high-dimensional nature of single cell cytometry data gives rise to numerous biologically relevant cell type hierarchies. treekoR acknowledges this by providing a framework which is not restricted to one specific cell type hierarchy constructed by a specific algorithm. We compared trees constructed using Hierarchical Ordered Partitioning And Collapsing Hybrid (HOPACH)^17^ clustering, average-linkage hierarchical clustering, and single-linkage hierarchical clustering via ‘tanglegrams’ (a pair of trees drawn with edges connecting matching leaves between the pair) on a PBMC sample from a healthy bone marrow donor^38^ (**Supplementary Figure 1b–1d**). The comparison highlights distinct cell type trees, which would consequently result in distinct quantifications of %parent proportions. Average-linkage hierarchical clustering - used in algorithms such as treeclimbr and citrus - and single-linkage clustering - closely resembling minimum spanning trees^39^ used in SPADE and visualising FlowSOM - generate distinct hierarchical representations of the data. In this dataset, HOPACH provided a representation more closely resembling the manual gating tree constructed by Bendall et al. Of importance is that one representation of cell type hierarchy may not necessarily be the most informative, but each of these representations can lead to diverse yet relevant %parent relationships.

When comparing the HOPACH constructed tree to the manual gating hierarchy, a clear difference is how NK cells and GMP cells are grouped - NK and GMP fall under the same immediate parent node in the HOPACH tree whilst they are not in the gating tree (**Supplementary Figure 1b**). An analyst may gate the cell types differently using two markers such as CD34 & CD45RA (**Supplementary Figure 1f**). Although this provides a more interpretable representation to cell groupings, it is clear in the t-SNE plot (**Supplementary Figure 1g**) how they would group together via clustering which considers all the available markers in unison. This indicates the effect of the experimental panel design, where some biologically distinct cell subpopulations can group together in automated clustering methods when there are more markers to distinguish them. Although there are notable differences between the automatically generated tree and the manually gated hierarchy, the HOPACH clustering is able to regenerate some of the cell type groupings. Despite some scenarios where the parent proportions may make less sense, this provides our framework with a big advantage over manual gating through being more efficient in handling larger datasets as well requiring a lower extent of prior knowledge to subset cells.

---

**Supplementary Fig 1:**
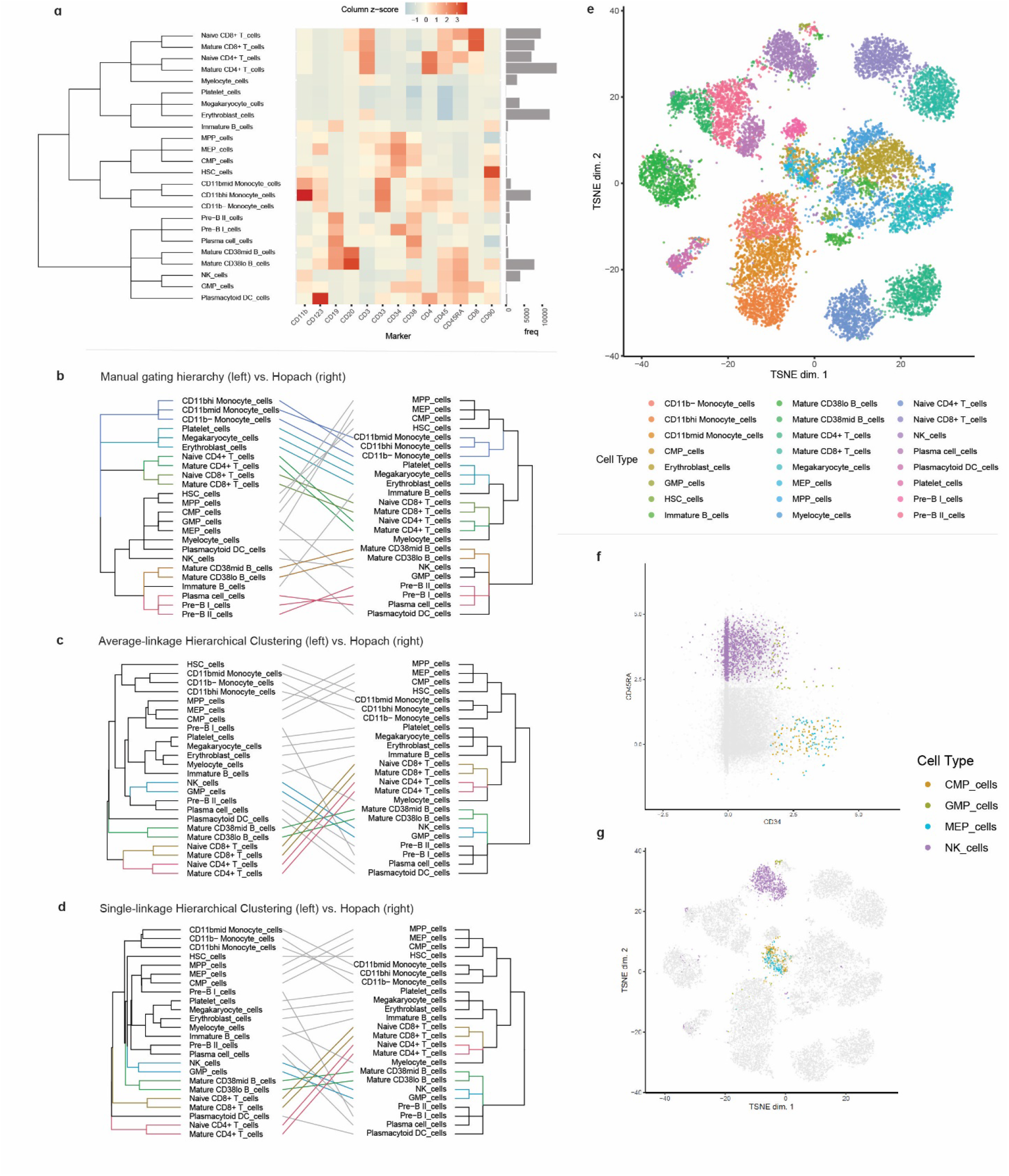
treekoR facilities varying cell type hierarchies to enable the measurement of cell types %parent. **a.** A heatmap of the median marker frequencies, clustered using HOPACH, for each cell type as subsetted with manual gating from a healthy human bone marrow sample^38^ (top left). **b.**Tanglegrams comparing two trees to highlight differences in cell type hierarchies: manual gating hierarchy vs. HOPACH clustering; **c.** average-linkage hierarchical clustering vs. HOPACH clustering; and **d.** single-linkage hierarchical clustering vs. HOPACH clustering (bottom right). **e.** A t-SNE plot of the sample, highlighted by the manually gated cell types **f.** Scatterplot of CD34 vs. CD45RA with only CMP, GMP, MEP & NK cells highlighted **g.** A t-SNE plot of the sample with only CMP, GMP, MEP & NK cells highlighted

## Notes

### Competing Interest Statement

The authors have declared no competing interest.

### Summary of Updates

Figure resolution updated; declarations updated; some grammatical nuances updated

